# Sleep deprivation impairs emotional adaptation to prolonged and ambiguous threat

**DOI:** 10.64898/2026.06.21.733648

**Authors:** Emma C. Sullivan, Cade McCall, Maximilian Croissant, Lisa-Marie Henderson, Guy Schofield, Scott A. Cairney

**Affiliations:** Department of Psychology, University of York, York, YO10 5DD, United Kingdom; Melbourne School of Psychological Sciences, University of Melbourne, Melbourne, VIC 3010, Australia; Department of Computer Science, University of York, York, YO10 5DD, United Kingdom; Institute of Mental Health Research, University of York, YO10 5DD, United Kingdom; Department of Archaeology, University of York, York, YO10 5DD, United Kingdom

## Abstract

Sleep problems are strongly linked to anxiety, but the psychological mechanisms underpinning this relationship are poorly understood. In this pre-registered study, we combined virtual reality, psychophysiology and multidimensional experience sampling to test the hypothesis that sleep deprivation disrupts emotional adaptation to sustained or resolving threat, giving rise to a heightened state of anxiety. Following a night of restful sleep or total sleep deprivation, healthy young adults navigated an immersive virtual world that alternated between ambiguously threatening and non-threatening contexts. Despite initial increases in emotional arousal, sleep-rested individuals quickly downregulated affective responses to ambiguous threat, reflecting efficient adaptation to the aversive virtual environment. Sleep-deprived individuals, by contrast, were unable to overturn elevated arousal responses, and exhibited a breakdown of goal-orientated, emotional control. Interestingly, resting heart rate variability, an index of affective regulatory capacity, mitigated arousal responses to ambiguous threat after sleep deprivation. These findings suggest that impaired adaptation to prolonged threat may represent a key mechanistic pathway linking insufficient sleep to anxiety.

## Introduction

Sleep problems are robustly linked to anxiety. A recent meta-analysis of 5,875 patients showed that sleep disturbances are highly prevalent in generalised anxiety disorder (1). Relatedly, a longitudinal study of 19,273 patients with insomnia revealed an 8.8-fold risk of developing anxiety, as compared to matched controls without insomnia (2). In healthy adults, a single night of experimental sleep deprivation drastically increased self-reported anxiety scores, with a large proportion of these sleep-deprived individuals temporarily exceeding the clinical threshold (3). Given the growing global burden of anxiety (4), understanding the affective and cognitive mechanisms underpinning the anxiogenic impacts of sleep deprivation is essential for developing targeted treatments and mitigation strategies.

One pathway through which sleep problems likely contribute to anxiety is via disruption of the brain’s threat processing networks. In healthy adults, a single night of sleep deprivation amplifies autonomic arousal responses to emotionally negative images (5). Similarly, functional neuroimaging studies have revealed a breakdown of top-down emotional control of amygdala by prefrontal cortex in sleep-deprived healthy adults exposed to aversive images and videos (3, 6); a brain activity profile that closely resembles that of clinically anxious individuals engaging in negative emotion processing (7–9). Moreover, sleep deprivation impairs reciprocal associations between central and peripheral emotion-signalling systems, disrupting accurate discrimination of threatening from non-threatening faces (10).

While these and other studies have provided important insights into the impact of sleep deprivation on emotional responses to threat, they have focused only on discrete and unambiguous events at fixed points in time (e.g., static images or short videos lasting only a few seconds). In real-world contexts, by contrast, the nature of threats can be uncertain even when an individual is aware of being at risk (11). Potential dangers are often more ambiguous and protracted than stimuli presented in laboratory settings, unfolding over several minutes and interspersed with periods of relative safety. Affective responses to prolonged and ambiguous threat can thus be adapted to the particular features of an experience and the familiarity that an individual has with their surroundings (12, 13). Determining how insufficient sleep influences emotional adaptation to sustained or resolving threat is therefore central to understanding how sleep problems give rise to anxiety.

We addressed this question in the current pre-registered study (osf.io/g4fte), combining virtual reality, psychophysiology and multi-dimensional experience sampling in N=54 healthy young adults. Participants underwent a night of restful sleep (with polysomnography) or total sleep deprivation before navigating an immersive virtual world that alternated between blocks of ambiguously threatening and non-threatening contexts (**Fig 1**). The ambiguously threatening contexts comprised dark, unsettling scenarios (e.g., an abandoned hospital ward) that were designed to evoke a sustained state of hypervigilance and anxiety, rather than the phasic arousal responses elicited by brief, unambiguous threats (11). The non-threatening contexts consisted of well-lit and benign office spaces. Skin conductance level (SCL) and heart rate (beats-per-minute, BPM) were monitored throughout the virtual experience to serve as real-time physiological indices of emotional arousal. Self-reported feelings of arousal were acquired retrospectively in a playback task. A video demonstration of the virtual world is available online (https://youtu.be/zm6pLPliZv0).

**Fig 1.**
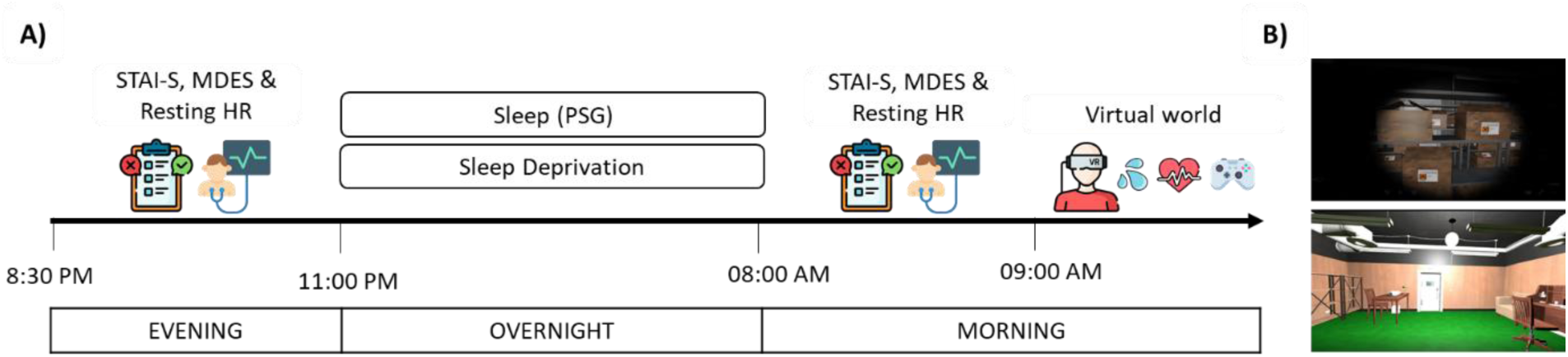
Experimental procedure and virtual world. **A)** Participants completed the State-Trait Anxiety Inventory—State Version (STAI-S), estimated the extent to which their ongoing thoughts matched a series of thought probes (Multidimensional Experience Sampling; MDES) and completed a 5-min resting heart rate recording before and after a night of restful sleep or sleep deprivation. Sleep was monitored with polysomnography (PSG), permitting a reliable assessment of slow-wave activity during non-rapid eye movement sleep. In the morning, participants entered an immersive virtual world while their skin conductance level and heart rate were monitored. Afterwards, they watched a replay of their virtual experience and retrospectively rated their feelings of emotional arousal (using a joystick). **B)** The virtual world alternated between blocks of ambiguously threatening (e.g., a storage room containing hazardous chemicals, top) and non-threatening scenarios (e.g., an office, bottom). There were two threatening and two non-threatening blocks in total, each lasting ∼2 min. Participants began in a well-lit office room that served as a non-threatening baseline.

We tested two hypotheses concerning the impact of sleep deprivation on emotional adaptation to prolonged, ambiguous threat. Our first hypothesis was that sleep loss would amplify emotional arousal responses during the ambiguously threatening blocks of the virtual world. Our second hypothesis was that sleep loss would slow the resolution of emotional arousal following transitions into non-threatening blocks of the virtual world, consistent with a failure to downregulate emotional affect once the potential danger has expired.

Given these proposed negative impacts of sleep deprivation, a converse question concerns the neurobiological mechanisms through which restful sleep prevents maladaptive emotional responding to evolving threats. Non-rapid eye movement slow-wave sleep and associated slow-wave activity (SWA, ∼0.5-4.0 Hz) have been implicated in anxiety regulation via the restoration of prefrontal emotional control networks (3). This aligns with disruptions to slow-wave sleep and SWA observed in patients with generalised anxiety disorder, post-traumatic stress disorder and insomnia, as well as subclinical populations with high trait anxiety (14–17). Accordingly, we further hypothesised that SWA during restful sleep would support emotional adaptation to evolving, ambiguous threat.

Successfully adapting emotional responses to changes in the external environment relies on flexible, goal-orientated cognition and the regulation of perseverative thoughts. Recent work has shown that sleep deprivation gives rise to intrusive, uncontrollable thoughts via a breakdown of prefrontal control and aberrant connectivity between brain networks supporting internally and externally focused mental processing (18). This raises the question of whether failures of emotional adaptation after sleep loss reflect disruptions of goal-orientated cognition and affective control, giving way to the repetitive and maladaptive thought patterns that reinforce anxiety. To explore this possibility, we had participants report on the content of their ongoing thoughts at rest and examined changes in latent thought patterns across restful sleep and sleep deprivation.

Finally, given sleep disturbance is a core feature of generalized anxiety disorder (19–21), identifying targets that could offset the negative impact of sleep loss on emotion regulation is essential for advancing treatment and mitigation efforts. Resting heart rate variability (HRV; the moment-to-moment variation between heartbeats) has been linked to flexible emotion processing (22) and cognitive control (23, 24). In earlier work, higher resting HRV was associated with improved regulation of emotional arousal in threatening virtual environments (25) and may support affective adaptation when sleep loss compromises regulatory capacity. We obtained resting heart rate recordings from each of our participants to explore this possibility.

## Materials and methods

### Pre-registration

The study methods and analyses were pre-registered (osf.io/g4fte). Deviations from our pre-registered methods are outlined and justified in Table S1.

### Participants

Eighty-five healthy young adults (aged 18-30 years) were recruited via an online recruitment platform and reported no history of neurological, psychiatric, attention, or sleep disorders. Participants completed several screening questionnaires and were excluded if they exceeded the clinical threshold on the Beck Depression Index (BDI-II ≥ 18; 26) and/or the Beck Anxiety Inventory (BAI ≥ 16; 27), if they had an extreme diurnal preference on the Morningness-Eveningness Questionnaire (MEQ < 31 or > 69; 28), or if they scored poorly on the Pittsburgh Sleep Quality Index (PSQI >6; 29). Inclusion was also contingent on participants reporting a consistent sleep schedule where they typically went to bed before 2am and rose by 8am having obtained at least six hours of sleep. Participants were excluded if they showed any abnormalities in physiological recordings (skin conductance levels and heart rate) during the virtual reality demo at session one (see Procedure). Participants were not using any prescribed medications except for the female contraceptive pill. Following standard procedures in our laboratory (30–34), participants were asked to refrain from caffeine and alcohol for 24 and 48 hours, respectively, before the study commenced and until the study ended.

Sixty-four participants met the inclusion criteria and completed the experimental protocol (sleep deprivation condition: N=36; sleep rested condition: N=28). N=8 participants were excluded from the sleep deprivation condition because they slept for >2 hours during the overnight interval, as indicated by self-report and/or actigraphy data, and another withdrew after feeling nauseated by the virtual environment. One participant was excluded from the sleep rested condition due to sleeping <4 hours during the overnight interval (as indicated by their polysomnography data). Therefore, our final sample size was N=54 (sleep deprivation condition: N=27, 18 female, mean ±SD age=19.59 ± 2.06 years; sleep rested condition: N=27, 15 female, mean ±SD age=20.30 ± 2.27 years). Participants in each condition were matched on trait anxiety levels as indicated by the State-Trait Anxiety Inventory—Trait Version (STAI-T; t = 0.45, *p* = .652; 34). They received £90 or BSc Psychology experimental participation credit for participating in the study. Ethical approval for this study was obtained by the Department of Psychology Research Ethics Committee at the University of York and all participants provided written informed consent.

Sample size requirements were informed by a power analysis. Our effects of interest were obtained from Ben Simon et al (3) who observed a deleterious effect of sleep deprivation on state anxiety, reflected by a significant interaction between condition (sleep rested or sleep deprived) and session (morning or evening; ɳ_ρ_^2^ = 0.36) and a positive correlation between SWA and state anxiety (*r* = .53). Our power analysis was based on the latter, smaller effect. This indicated that N=27 participants were required to detect an effect of r = .53 (90% power, α = 0.05, one-tailed) in the sleep rested condition.

### Procedure

The study protocol comprised three sessions. In session one, participants completed an online screening questionnaire and a short virtual reality demo. This demo ensured that participants could acclimatise to the controls without being exposed to any ambiguous threat. Skin conductance levels (SCLs) and heart rate were recorded to mimic the main virtual environment and screen for any SCL-non-responders or electrocardiography (ECG) irregularities. Participants completed session one at least 24 hours before session two.

At 9am on the morning of session two, participants collected an actigraphy watch and were instructed to wear it until the end of the study. During session two (∼8.30pm), they underwent a 5-min resting heart rate recording and then completed the multidimensional experience sampling (MDES) protocol (36–40). For MDES, participants were presented with a series of 13 thought probes and, for each, were asked to indicate on a scale from 1 (Not at all) to 10 (completely), the extent to which their recent thoughts were aligned with the probe (see Table S2 for the full set of thought probes). Afterwards, participants completed the State-Trait Anxiety Inventory—State Version (STAI-S; 34), the Stanford Sleepiness Scale (SSS; 40) and the psychomotor vigilance task PVT; 41). They were then informed whether they had been assigned to the sleep rested or sleep deprivation condition.

Sessions two and three were separated by an overnight interval during which participants either remained awake (i.e., were sleep deprived) or slept in the sleep laboratory. Participants in the sleep deprivation condition were sent home and permitted to communicate, read, use electronic devices, watch TV, or play games^1^. Adherence to the sleep deprivation protocol was confirmed with actigraphy, which verified that all sleep-deprived participants remained awake throughout the night. They also completed an electronic questionnaire every 30 minutes between 11pm and 6.30am, with 95.48% of items answered before returning to the lab at 8am^2^. In the sleep-rested condition, participants were wired up for polysomnography and lights were turned out at ∼11pm. They were woken up at ∼7am.

Session three began at ∼8.30am and participants repeated the MDES protocol, questionnaires and PVT. Sleep-deprived participants also completed a questionnaire probing their adherence to the sleep deprivation protocol and activities they engaged in throughout the night (see Table S3 for further details). Participants then entered the immersive virtual world while SCL and heart rate were continuously monitored.

The virtual world began with an auditory prelude that included a dramatic background story and task instructions. Participants were told that they had woken in a laboratory after a violent incident, where researchers were being held captive by a malevolent group seeking to destroy an experimental drug designed to eradicate evil. The researchers’ survival depended on them collecting red and blue lanterns. Participants first started the virtual task in a well-lit office room, with standard furnishings and gentle music, which served as a non-threatening baseline. After 1 minute, a freight elevator door opened and participants descended to a basement level where they moved through the first block of ambiguously threatening rooms (Hospital Ward, Storage Room, Autopsy Room). These rooms were designed to evoke sustained unease rather than discrete unambiguous scares, featuring dark, unsettling environments requiring torchlight, ominous music and menacing sound effects (e.g., footsteps, whispers; 11, 42–45). Participants spent 30 seconds in each room during which they were instructed to explore and retrieve the lanterns before the door to the next room opened. This block was followed by a non-threatening block of three separate office spaces, similar to the initial office room, with identical instructions. Participants then completed a second ambiguously threatening block (Bathroom, Torture Chamber, Kitchen) before ascending via the freight elevator back to an office room, similar to the one that they had started in, for 30 seconds (serving as a second non-threatening block). A video of the virtual world is available at https://youtu.be/zm6pLPliZv0.

Next, participants completed a playback task where they watched a first-person recording of their virtual experience and were instructed to retrospectively rate how they felt during every moment of the experience, using a joystick to position a curser on an affect grid with valence (unpleasant-pleasant) and arousal (excited-not at all excited) on the x axis and y axis, respectively (47, 48). The magnitude of each axis varied from −100 to 100 and a circle depicted the current position of the joystick. Only responses on the arousal dimension of the affect grid were used in our analyses.

After completing the playback task, participants returned the actigraphy watch and were debriefed before leaving the laboratory.

### Materials and equipment

#### Questionnaires

Self-reported anxiety was measured with the State-Trait Anxiety Inventory—State Version (35). This comprises 20 items and indexes an individual’s feelings of anxiety at the current moment. Responses on each item vary from 1 (not at all) to 4 (very much so). Total state anxiety scores range from 20–80, with higher scores reflecting higher levels of state anxiety.

Sleepiness was measured with the Stanford Sleepiness Scale (41). Respondents choose one of seven statements that best describes their current state, with each statement corresponding to an increasing level of sleepiness.

#### Psychomotor vigilance task

Participants were instructed to click a mouse button as quickly as possible when a digital counter appeared on the screen (42). The inter-stimulus interval (ISI) varied from 2 to 10 s and task duration is ∼3 min. Response time feedback was provided on each trial.

#### Heart rate variability (HRV)

5-minute resting heart rate recordings were administered before and after a night of sleep or sleep deprivation. We calculated the root mean square of successive difference (RMSSD) between normal heartbeats (a measure of vagally mediated HRV), as it is the primary time-domain measure to estimate HRV (49) and has been shown to be reliable for short recordings (50). Participants were instructed to relax and remain still for 7 minutes without closing their eyes or crossing their feet. The first 2 minutes of data were excluded to permit acclimatisation. For details about equipment and data processing, see Methodological Details S1.

#### Virtual world

The virtual world was adapted from the *Underwood Project*: a modular virtual environment for eliciting ambiguous threat, built in the Unity game creation environment (11). Participants experienced the virtual world through an HTC Vive head-mounted display unit with an integrated Dual AMOLED 3.6-inch diagonal screen (resolution: 1080 × 1200 pixels per eye, refresh rate: 90 Hz, field of view: 110°). A wireless Vive controller was used so that participants could move around the virtual world using the dual-stage trigger. Audio was played through DOQUAS wireless headphones from a standard desktop computer.

SCL and heart rate in the virtual world were recorded, z-scored within person and aggregated across the threatening and non-threatening blocks of the virtual world (i.e., separately for the first threatening block, first non-threatening block, second threatening block and second non-threatening block). For details about equipment and data processing, see Methodological Details S2.

The playback task was administered on a standard desktop computer using the open-source software DARMA (47). Participants rated how they felt during every moment using the joystick of a wireless controller (Microsoft Xbox). Subjective arousal ratings from the playback task were exported into R studio and z-scored within person, before being aggregated across the threatening and non-threatening blocks of the virtual world.

#### Sleep monitoring

Polysomnographic data was recorded to ensure that participants in the sleep group obtained an adequate amount of sleep and allowed us to measure whether SWA supports emotional adaptation to evolving ambiguous threat. For details about equipment and data processing, see Methodological Details S3.

Actigraphy data were acquired with a Philips Actiwatch 2 (Philips Respironics), which participants wore throughout the study.

### Analysis

Unless otherwise stated, all analyses were run in R version 4.5.2 (51), and all figures were created using the *ggplot2* package (52).

To examine whether sleep deprivation amplified emotional responses to ambiguous threat, we first calculated arousal-change scores from baseline (the office room at the beginning of the virtual world) during the first and second threatening blocks. For each participant, aggregated baseline scores were subtracted from aggregated scores in each threatening block, separately for SCL, heart rate and subjective arousal. These difference scores indexed the change in arousal from baseline in within-person standard deviation units, with higher scores indicating greater increases in arousal.

We also calculated change scores to investigate how sleep deprivation affected the resolution of emotional arousal once the ambiguous threat had subsided. For each measure (SCL, heart rate and subjective arousal), aggregated values for the first and second threatening blocks were subtracted from the aggregated values for the adjacent first and second non-threatening blocks. Lower scores indicated a greater reduction in emotional arousal (in standard deviation units) upon reaching a non-threatening block.

Change scores were applied to two-way mixed ANOVAs with the between-subjects factor *Group* (Sleep Rested or Sleep Deprived) and the within-subjects factor *Block* (One or Two). Where significant *Group*Block* interactions were significant, Bonferroni-corrected pairwise comparisons were performed. Partial eta squared was reported for all ANOVAs and Cohen’s d for all pairwise tests. These analyses were conducted using the *rstatix* package (53). To assess whether SWA supports emotional adaptation to ambiguous threat, we calculated change scores reflecting the difference in emotional arousal between the first and second threatening blocks of the virtual world (block two minus block one), for SCL, HR and subjective responses. Lower scores indicated a greater reduction in emotional affect across blocks. These change scores were then correlated with SWA in the sleep rested condition. Bonferroni-corrected p-values were used to account for multiple comparisons, all correlations were analysed using the *Hmisc* package (54).

To explore whether emotional dysregulation after sleep deprivation was related to impairments of goal-orientated cognition, multidimensional experience sampling scores were entered into a principal components analysis (PCA) using the *psych* package (55). In keeping with earlier work (18), responses to the 13 thought probes from the evening and morning sessions were concatenated into a single matrix, and PCA with varimax rotation was performed. MDES data were z-scored prior to analysis. Components were identified based on eigenvalues, and the inflection point in the scree plot (see Figure S1). Component scores were extracted for each participant and applied to a two-way mixed ANOVA with the between-subjects factor *Group* (Sleep Rested or Sleep Deprived) and the within-subjects factor *Session* (Evening or Morning). For significant *Group*Session* interactions, Bonferroni-corrected pairwise comparisons were performed. Fourteen participants were excluded from these analyses due to missing item data at one or both sessions.

To examine whether higher levels of resting HRV mitigated the impact of sleep deprivation on adaptation to ambiguous threat, participants were stratified into low and high HRV groups, using a median split of RMSSD from the evening session (56). Change scores for self-reported arousal and SCLs were applied to a three-way mixed ANOVAs with the between-subject factors *Group* (Sleep Rested or Sleep Deprived) and *HRV (*Low or High), and the within-subjects factor *Block* (Threatening Block One or Two). As in the main analysis, when interactions involving HRV (e.g., *Group*HRV, Block*HRV, Group*Block*HRV*) were significant, Bonferroni-corrected pairwise comparisons were performed.

## Results

### Sleep deprivation escalates anxiety

To confirm that sleep deprivation elevated self-reported feelings of anxiety, we first examined overnight changes on the State-Trait Anxiety Inventory—State Version (STAI-S; 26, **Fig 1A**). Whereas STAI-S scores decreased from evening to morning in sleep-rested individuals (t = 2.74, *p* = .011, d = 0.53), they increased markedly in sleep-deprived individuals (t = 6.25, *p* < .001, d = 1.20; F(1, 52) = 45.28, *p* < .001, ɳ_ρ_ ^2^ = 0.47; **Fig 2A**). Notably, in the morning after sleep deprivation, 56% of participants (N=15) exceeded the STAI-S threshold for clinically significant anxiety, whereas only 7% (N=2) participants surpassed this threshold following restful sleep. No significant between-group difference was observed at the evening baseline session (t = 0.27, *p* = .788, d = 0.07). These findings replicate previous work demonstrating that a single night of sleep deprivation leads to an elevated state of anxiety (3, 57).

**Fig 2.**
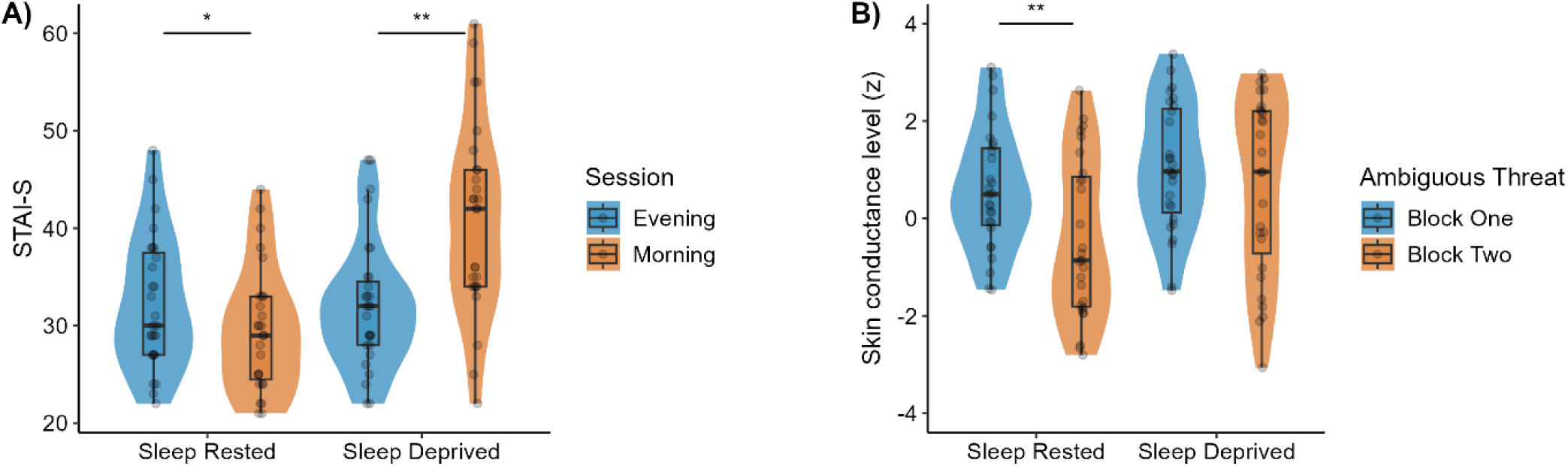
Sleep deprivation impairs emotional adaptation to ambiguous threat. **A)** State anxiety decreased overnight in sleep-rested individuals but increased markedly overnight in sleep-deprived individuals (based on the State-Trait Anxiety Inventory – State Version; STAI-S). **B)** Skin conductance levels (SCLs) decreased between the first and second threatening blocks of the virtual world in sleep-rested individuals but remained elevated in sleep-deprived individuals. Violin plots: the width of the shaded area represents the proportion of data located there. Boxplots depict the minimum, first quartile, median, third quartile, and maximum values. Individual data points are marked in grey. ** p < .001, * p < .05.

Alertness levels were reduced after sleep deprivation relative to restful sleep, as indicated by the Stanford Sleepiness Scale (28, t = 8.16, *p* < .001, d = 2.22) and psychomotor vigilance task (29, t = 2.58, *p* = .013, d = 0.70). Extended alertness data are available in Note S1 and Table S4.

### Sleep loss prevents emotional adaptation to ambiguous threat

We then turned to the impacts of sleep deprivation on emotional adaptation to prolonged and ambiguous threat. To determine whether sleep loss amplified emotional responses to ambiguous threat, we assessed arousal levels during the first and second threatening blocks of the virtual world (**Fig 1B**). SCLs increased relative to baseline across all participants during the first threatening block (F(1,52) = 24.75, *p* < .001, ɳ_ρ_^2^ = 0.32). Importantly, whereas SCLs decreased during the second threatening block in sleep-rested participants (t = 5.58, *p* < .001, d = 1.07), they remained elevated in sleep-deprived participants (t = 1.80, *p* = .084, d = 0.35, F(1,52) = 4.98, *p* = .030, ɳ_ρ_^2^ = 0.09; **Fig 2B**), suggesting that sleep loss prevented a recalibration of emotional responses to sustained ambiguous threat. SCLs did not differ significantly between groups when averaged across blocks (F(1,52) = 3.95, *p* = .052, ɳ_ρ_^2^ = 0.07). Parallel analyses of heart rate (BPM) and self-reported feelings of arousal (assessed retrospectively via a playback task) did not reveal significant differences between groups (see Note S2 and Table S5).

To establish whether sleep deprivation obstructed the resolution of emotional reactivity once the ambiguous threat had dissipated, we assessed changes in arousal between the threatening and non-threatening blocks of the virtual world. Across all participants, SCLs decreased following transitions into non-threatening blocks, with a larger decrease observed during the first than the second transition (F(1,52) = 7.83, *p* = .007, ɳ_ρ_^2^ = 0.13). However, sleep deprivation had no significant impact on the magnitude of this change (*Group*: F(1,52) = 0.72, *p* = .400, ɳ_ρ_^2^ = 0.01, *Group*Block*: F(1,52) = 2.98, *p* = .090, ɳ_ρ_^2^ = 0.05). Again, no significant between-group differences were observed in parallel analyses of heart rate and self-reported feelings of arousal (see Note S3 and Table S6).

### Emotional adaptation is not associated with slow-wave activity

We next investigated whether SWA (0.8–4.6 Hz) during non-rapid eye movement sleep supported emotional adaptation to prolonged, ambiguous threat. No significant correlations were observed between SWA and either overnight reductions in state anxiety (STAI-S: r = .09, *p* = 1.00) or the attenuation of SCLs between the first and second blocks of ambiguous threat (r = −.08, *p* = 1.00). Correlations between SWA and our other measures of emotional arousal were also not significant (see Table S7).

### Lack of sleep disrupts goal-orientated cognition

We then explored whether maladaptive emotional responding after sleep deprivation was accompanied by a breakdown of the top-down control processes required to regulate emotions effectively. We used multidimensional experience sampling (MDES)—an established thought sampling technique (36–40) where individuals rate the extent to which their ongoing thoughts correspond to a set of thought probes (e.g., *my thoughts involved future events*)— in the evening before restful sleep or sleep deprivation, and again in the morning. Principal components analysis was then applied to the MDES data to identify latent dimensions of thought that capture shared variance.

The first identified component captured a *goal orientated* dimension corresponding to deliberate future problem solving. Interestingly, while there was no significant between-group difference in goal orientated thinking at the evening session (t = 0.72, *p* = .474, d = 0.23), one did emerge the following morning, with sleep deprived participants engaging in fewer goal-orientated thoughts than the sleep-rested participants (t = 2.66, *p* = .012, d = 0.85; F(1,38) = 4.07, *p* = .051, ɳ_ρ_^2^ = 0.10; **Fig 3A**). Sleep deprivation also resulted in significantly less goal-orientated thinking overall (F(1, 38) = 4.88, *p* = .033, ɳ_ρ_^2^ = 0.11). The general difference between sessions was not significant (F(1, 38) = 1.22, *p* = .276, ɳ_ρ_^2^ = 0.03).

**Fig 3.**
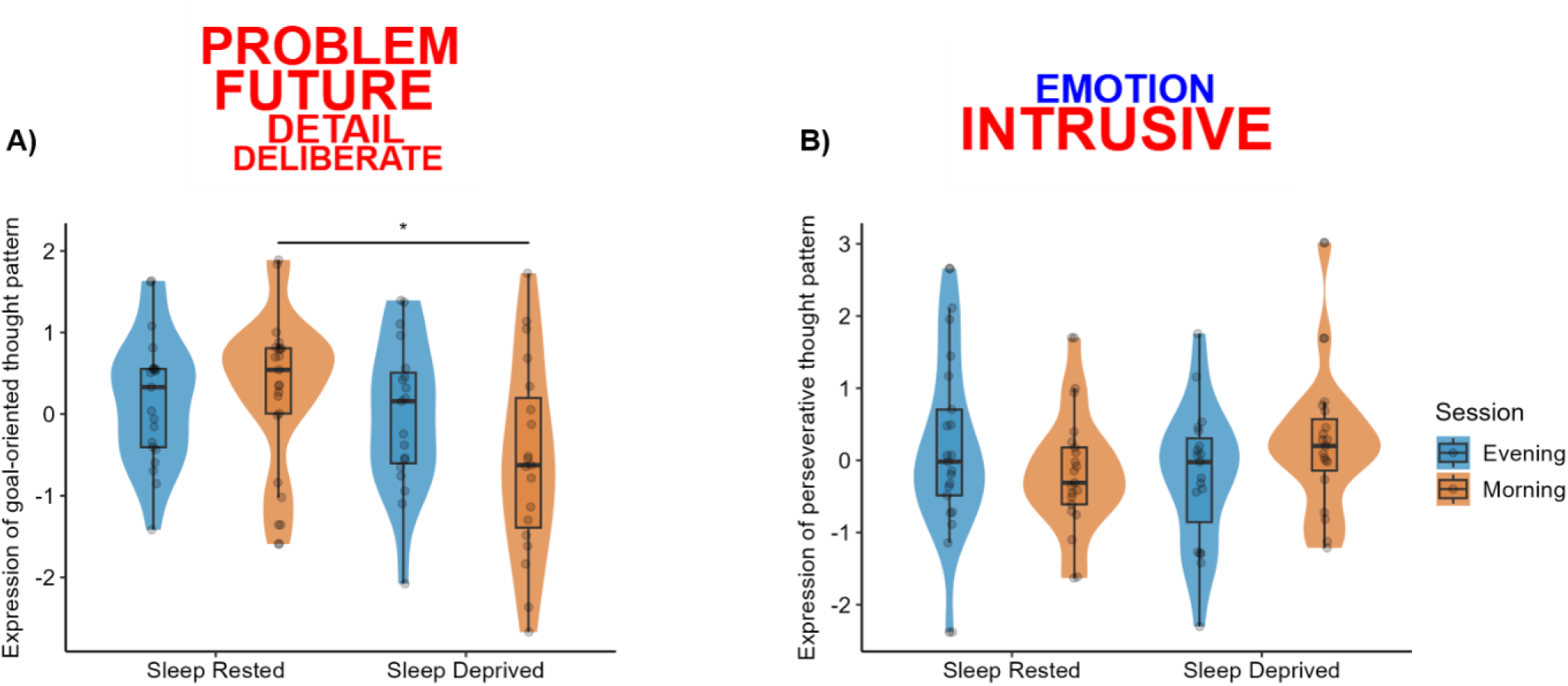
Sleep deprivation disrupts top-down control. Principal components analysis of multidimensional experience-sampling data was used to identify latent patterns of thought by grouping thought probes that capture shared variance. **A)** The first component corresponded to a pattern of goal-orientated thinking. Component loadings are presented as a word cloud, where the word size reflects loading magnitude and colour indicates direction of the relationship (red: positive, blue: negative). Sleep deprived participants reported fewer goal-orientated thoughts than the sleep-rested participants in the morning session, but not in the evening session. **B)** The second component reflected a perseverative dimension of thought. Perseverative thinking increased after sleep deprivation compared to restful sleep, although this difference did not reach statistical significance. * p < .05.

The second component captured a *perseverative* dimension corresponding to intrusive and negative thoughts. Although overnight changes in perseverative thinking significantly differed between the sleep-rested and sleep-deprived participants (F(1,38) = 4.77, *p* = .035, ɳ_ρ_^2^ = 0.11), direct comparisons between these groups did not reach significance at the evening (t = 1.17, *p* = .251, d = 0.37) or morning sessions (t = -1.53, *p* = .135, d = 0.49; **Fig 3B**). No significant overall difference emerged between groups (F(1, 38) < 0.01, *p* = .943, ɳ_ρ_^2^ < 0.01) or sessions (F(1,38) = 0.05, *p* = .830, ɳ_ρ_^2^ < 0.01). See Figure S1 for analysis of the other components.

### Heart rate variability mitigates affect dysregulation after sleep loss

Finally, we explored whether the negative impact of sleep deprivation on emotional adaptation was mitigated by higher levels of heart rate variability (HRV) at rest (based on a 5-min resting HR recording acquired in the evening before restful sleep or sleep deprivation), a putative index of flexible emotion regulation and inhibitory control (22–25).

Participants with low HRV (based on a median split) had significantly higher self-reported levels of arousal during the ambiguously threatening blocks of the virtual world after sleep deprivation than restful sleep (t = -4.83, *p* < .001, d = 1.34). Contrastingly, no significant difference emerged in the high HRV group (t = 1.22, *p* = .228, d= 0.34; F(1, 50) = 6.15, *p* = .017, ɳ_ρ_^2^ = 0.11; **Fig 4**). The mitigating impact of HRV on the maladaptive influences of sleep deprivation did not significantly differ between blocks (F(1, 50) = 0.01, *p* = .937, ɳ_ρ_^2^ < .01) and overall levels of self-reported arousal did not significantly differ between HRV groups (F(1, 50) = 0.00, *p* = .974, ɳ_ρ_^2^ < .01; *HRV*Block*: F(1, 50) = 0.05, *p* = .824, ɳ_ρ_^2^ < .01).

**Fig 4.**
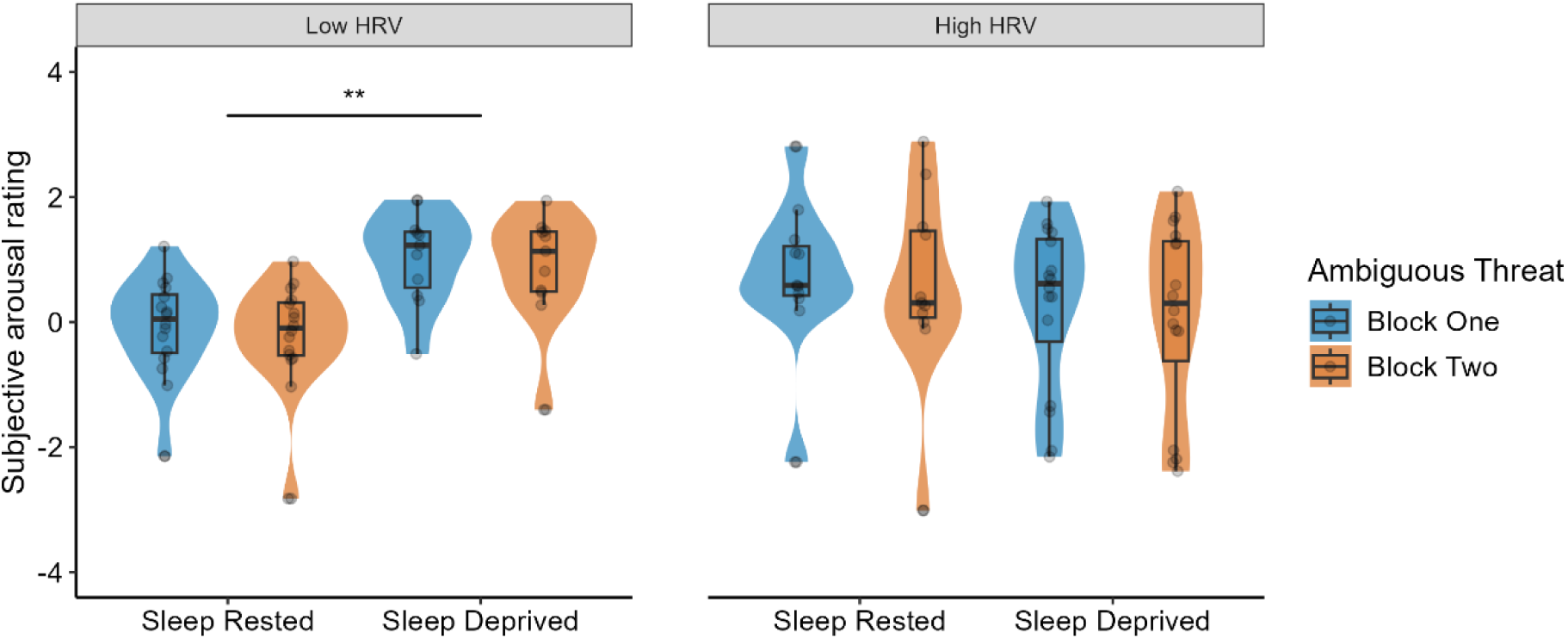
Heart rate variability mitigates the negative effect of sleep deprivation on emotional adaptation. Across both ambiguously threatening blocks of the virtual world, self-reported arousal was significantly higher following sleep deprivation compared to restful sleep in participants with low HRV, but not high HRV (root mean square of successive differences) ** p < .001.

The same analysis of SCL data did not produce any significant effects, suggesting that the protective effect of high HRV is expressed at the level of subjective experience, rather than in psychophysiological indices of arousal, consistent with our prior work (25).

## Discussion

We combined immersive virtual reality, psychophysiology, and multidimensional experience sampling in healthy young adults to examine how sleep deprivation affects emotional adaptation to sustained or resolving threat. Whereas sleep-rested individuals rapidly acclimatised to ambiguously threatening contexts, sleep-deprived individuals showed persistently elevated arousal responses to prolonged, ambiguous threat. However, changes in emotional arousal were comparable across the sleep-rested and sleep-deprived conditions when participants moved from threatening to non-threatening contexts. Sleep deprivation also decreased goal-orientated patterns of thought, consistent with a breakdown of top-down cognitive control. Interestingly, emotional adaptation was better preserved in sleep-deprived participants with higher resting heart rate variability (HRV), suggesting a moderating influence of HRV on the anxiogenic impacts of sleep loss. Together, these findings shed new light on the affective and cognitive pathways through which an absence of sleep gives rise to maladaptive patterns of emotional arousal and anxiety.

Before examining the impact of sleep deprivation on emotion adaptation in the context of an immersive virtual world, we first assessed whether a night of sleep deprivation altered participants’ general state of anxiety, as measured by the State-Trait Anxiety Inventory—State Version (STAI-S). Sleep-deprived participants exhibited a significant overnight increase in state anxiety, with 56% exceeding the STAI’s clinical threshold during the morning session (none of whom had done so prior to the overnight interval). In contrast, sleep-rested participants showed a significant reduction in state anxiety from evening to morning, with only 7% exceeding the clinical threshold. These findings replicate previous work in healthy young adults (3, 57) and are consistent with the high comorbidity between sleep disturbances and anxiety disorders in the general population (1, 2, 58).

Having established a generalised increase in state anxiety after sleep deprivation, we asked whether an absence of sleep impaired emotional adaptation to prolonged, ambiguous threat. Consistent with this view, elevated arousal responses (based on SCLs) persisted across the first and second blocks of ambiguous threat in the sleep deprivation condition but dissipated across blocks in the sleep rested condition. This suggests that sleep deprivation prevents the emotional brain from acclimatising to protracted and ambiguous threat, giving rise to an elevated state of anxiety.

It is worth noting that analogous effects were not observed in our heart rate or self-report data. This potentially reflects the greater specificity of SCL to changes in sympathetic arousal, relative to the integrated sympathetic and parasympathetic influences that shape heart rate fluctuations (59, 60). Furthermore, our self-report data relied on retrospective assessments that may have been less sensitive to sleep loss than electrodermal measures collected in real time.

Given the negative impact of sleep deprivation on the adaptation of emotional responses to prolonged, ambiguous threat, we expected that sleep loss would also disrupt the resolution of emotional arousal once the threat had dissipated. However, while transitioning between threatening and non-threatening contexts prompted a general decrease in SCLs, this reduction in autonomic arousal did not significantly differ across the sleep-deprived and sleep-rested conditions. One explanation for this outcome relates to the varied cognitive demands of our immersive virtual world. Although top-down control processes are likely required to navigate ambiguously threatening contexts (e.g., to downregulate negative emotions and unwanted thoughts), these processes become less necessary after the danger has passed. From this perspective, the adverse effects of sleep loss on emotional and cognitive control would be expected to manifest as a failure to acclimatise to prolonged, ambiguous threat, but not as impairments in affective recovery upon reaching points of relative safety.

As a corollary to this view, if sleep deprivation impairs the top-down control mechanisms responsible for adaptively regulating emotions when confronted with an ambiguous threat, then sleep-deprived individuals should engage in fewer goal-orientated thoughts than their sleep-rested counterparts. We explored this possibility by applying principal components analysis to our multidimensional experience sampling data. Whereas no between-condition differences were observed in the evening session, the sleep-deprived individuals reported fewer goal-directed than the sleep-rested participants in the morning session, consistent with a recent study using a similar approach (18).

Previous work has suggested that SWA during non-rapid eye movement sleep regulates anxiety via the restoration of prefrontal emotional control networks (3), raising the possibility that SWA also supports emotional adaptation when exposed to prolonged, ambiguous threat. However, SWA was not significantly correlated with either the overnight change in state anxiety or the change in SCLs across the first and second blocks of ambiguous threat in the immersive virtual world. These null results may be considered surprising given the reductions in SWA and non-rapid eye movement slow-wave sleep (where SWA predominates) observed in patients with anxiety related disorders and insomnia, as well as in non-clinical populations with high-trait anxiety (14–17), and highlight the need for further research to better understand the anxiolytic function of sleep.

We also sought to explore factors that may mitigate the impact of sleep deprivation on emotional adaptation, focusing on resting heart rate variability (HRV), given its putative role in affective and cognitive control (22–24). Sleep loss increased arousal responses to ambiguous threat in participants with low HRV, but not participants with high HRV, raising the possibility that high HRV buffers against the deleterious impacts of sleep deprivation on emotional flexibility. Interestingly, this moderating effect of HRV emerged in our self-reported arousal data (acquired via the playback task) and not in the SCL data, mirroring earlier work (25). This may suggest that the sympathetic–parasympathetic balance, as indexed by high HRV, is better reflected in subjective, experiential regulation than in SCL.

In conclusion, we show that sleep deprivation disrupts emotional adaptation to prolonged and ambiguous threat, arising from a breakdown of goal-orientated cognition and leading to a heightened state of anxiety. Higher HRV confers resilience to the adverse effects of sleep loss on emotional control and warrants further investigation as a potential translational target. These findings provide new mechanistic insight into the central role of sleep problems in the emergence, persistence and resolution of anxiety.

## Data availability

The data that support the findings of this study are available on the Open Science Framework (https://osf.io/m7tfd).

## Code availability

The code used to analyse the data presented in this study is available on the Open Science Framework (https://osf.io/m7tfd).

## Acknowledgements

This research was supported by the University of York Department of Psychology Doctoral Studentship (ECS) and Medical Research Council Career Development Award (MR-P020208-1; SAC). SAC is funded by the European Union (ERC, SLEEPAWAY, 101169737). Views and opinions expressed are however those of the authors only and do not necessarily reflect those of the European Union or the European Research Council. Neither the European Union nor the granting authority can be held responsible for them.

## Footnotes

1 The decision to send participants home was in response to an institutional recommendation to minimise face-to-face contact as the UK emerged from the COVID-19 pandemic.

2 Two sleep deprived participants did not complete the questionnaire due to technical difficulties. However, actigraphy data confirmed they remained awake throughout the night.

