## Supplementary Information for "Sleep deprivation impairs emotional adaptation to prolonged and ambiguous threat"

**Table S1.** List of deviations from the pre-registration and their justifications.

| Pre-registration | Manuscript |
|---|---|
| **1. Data preprocessing: SCL** | |
| SCLs will be square root transformed due to the possibility of a non-normal distribution. | SCL values were z-scored within-participants. |
| **Justification:** We changed our transformation for SCL values from square root transformation to z-scoring to maintain consistency with our other outcome measures (i.e., HR and self-reported arousal). | |
| **2. Data analysis: calculation of arousal change** | |
| Arousal-change scores will be calculated by subtracting mean arousal in block *n* from mean arousal in block *n*-1 | To calculate arousal-change scores from baseline, mean arousal at baseline was subtracted from mean arousal in each threatening block. |
| **Justification:** To examine whether sleep deprivation amplified emotional responses to ambiguous threat, it was deemed more appropriate to examine increases in arousal during the threatening contexts relative to a non-threatening baseline. | |
| **3. Data analysis: correlations between SWA and arousal** | |
| We will examine the correlation between SWA and mean arousal in the first threatening block. | We examined the correlation between SWA and the change in arousal between the first and second threatening blocks. |
| **Justification:** Given our hypothesis that SWA supports the resolution of emotional arousal to prolonged, ambiguous threat, it was deemed more appropriate to measure arousal change between the first and second threatening blocks. | |

**Table S2.** Multidimensional experience sampling (MDES) thought probes.

| Dimension | Description | 1 | 10 |
|---|---|---|---|
| Task | My thoughts were focused on the task: | Not at all | Completely |
| Future | My thoughts involved future events: | Not at all | Completely |
| Past | My thoughts involved past events: | Not at all | Completely |
| Self | My thoughts involved myself: | Not at all | Completely |
| Person | My thoughts involved other people: | Not at all | Completely |
| Emotion | The emotion of my thoughts was: | Negative | Positive |
| Modality | My thoughts were in the form of: | Images | Words |
| Detail | My thoughts were detailed and specific: | Not at all | Completely |
| Deliberate | My thoughts were: | Spontaneous | Deliberate |
| Problem | I was thinking about solutions to problems (or goals): | Not at all | Completely |
| Diverse | My thoughts were: | One topic | Many topics |
| Intrusive | My thoughts were intrusive: | Not at all | Completely |
| Source | My thoughts were linked to information from: | Environment | Memory |

**Table S3.** A list of activities that those in the sleep deprivation condition reported engaging at some stage during the overnight interval (N=27). These categories were based on self-report.

| Self-reported activity | % of participants who self-reported engaging in the activity |
|---|---|
| Watching TV | 88.89 |
| University work | 48.15 |
| Organising/cleaning | 44.44 |
| Exercising | 37.04 |
| Preparing/eating food | 37.04 |
| Gaming | 33.33 |
| Recreational activities | 29.63 |
| Browsing social media | 29.63 |
| Socialising with friends | 29.63 |
| Self-care | 18.52 |
| Other | 7.41 |

**Table S4.** Means and standard errors for alertness measures across sessions.

| Alertness measure | Sleep rested (N = 27) | Sleep deprived (N = 27) |
|---|---|---|
| **SSS** | | |
| Evening | 2.70 [*0.21*] | 2.59 [*0.20*] |
| Morning | 2.15 [*0.21*] | 4.93 [*0.27*] |
| **PVT** | | |
| Evening | 273.85 [*7.82*] | 272.51 [*5.22*] |
| Morning | 278.21 [8.*10*] | 308.76 [*8.65*] |

SSS = Stanford Sleepiness Scale (maximum score 7). PVT = Psychomotor vigilance task, as assessed by examining response times in milliseconds.

**Table S5.** Means and standard errors for the changes in skin conductance level (SCL), heart rate and subjective arousal ratings across the two ambiguously threatening blocks of the virtual world (relative to baseline; z-scores).

| Arousal measure | Sleep rested (N = 27) | Sleep deprived (N = 27) |
|---|---|---|
| **SCL** | | |
| Block One | 0.58 [*0.24*] | 1.05 [*0.25*] |
| Block Two | −0.42 [*0.31*] | 0.67 [*0.35*] |
| **Heart rate** | | |
| Block One | 0.38 [*0.11*] | 0.40 [*0.13*] |
| Block Two | 0.55 [*0.12*] | 0.50 [*0.11*] |
| **Self-reported arousal** | | |
| Block One | 0.25 [*0.20*] | 0.59 [*0.22*] |
| Block Two | 0.10 [*0.24*] | 0.40 [*0.26*] |

**Table S6.** Means and standard errors for skin conductance level (SCL), heart rate and subjective arousal ratings across the two transitions from threatening to non-threatening blocks of the virtual world (z scores).

| **Arousal measure** | **Sleep rested (N = 27)** | **Sleep deprived (N = 27)** |
|---|---|---|
| **SCL** | | |
| Block One | −0.88 [*0.15*] | −0.50 [*0.15*] |
| Block Two | −0.26 [*0.15*] | −0.36 [*0.16*] |
| **Heart rate** | | |
| Block One | 0.04 [*0.10*] | 0.01 [*0.10*] |
| Block Two | 0.20 [*0.12*] | 0.11 [*0.10*] |
| **Self-reported arousal** | | |
| Block One | −1.05 [*0.18*] | −0.82 [*0.22*] |
| Block Two | −1.19 [*0.27*] | −0.94 [*0.28*] |

**Table S7.** Pearsons correlations between slow-wave activity (SWA) and attenuation of emotional arousal (heart rate, self-reported arousal ratings) between the two ambiguously threatening blocks of the virtual world.

| Arousal measure | Total SWA (% power) |
|---|---|
| Heart rate | −.23 [−.56-.16] |
| Self-reported arousal* | .33 [−.10-.68] |

Values in square brackets indicate the 95% confidence interval for each correlation. All *p-values* were Bonferroni corrected and were > .05. *Spearman correlation. The confidence intervals of the Spearman's rank correlation coefficients were computed by bootstrapping.

**Methodological Details S1.** Heart rate variability.

Resting heart rate was recorded using a BIOPAC MP36R data acquisition system with three-lead ECG and disposable Ag/AGCl foam electrodes (BIOPAC, EL503). Electrodes were placed on the sternal end of the right clavicle, left mid-clavicle (ground electrode), and lower left rib cage. Data were recorded via AcqKnowledge software (version 4.4.1).

R-peaks (the maximum amplitude of an R wave in a QRS complex) were first identified using an automated routine in Acqknowledge. Missed and misclassified R-peaks were then manually added and removed, respectively. The interbeat interval time series was then exported into Kubios HRV Standard 3.5.0 Software (1).

**Methodological Details S2.** Skin conductance level (SCL) and heart rate.

Psychophysiological signals in the virtual world were recorded using a BIOPAC MP160 acquisition system with AcqKnowledge software and sampled at 2000 Hz. SCL was recorded using a wireless BIOPAC BioNomadix amplifier (BN-PPGED) with a BioNomadix dual electrode lead and disposable Ag/AgCl foam electrodes (BIOPAC, EL507a). Electrodes were attached to the middle phalanges of the left middle and index fingers using an isotonic electrode paste (BIOPAC Gel 101a) and the BioNomadix transmitter was placed on participant's nondominant wrist. The system clocks of the psychophysiological acquisition computer and the rendering computer for the virtual world were synchronised to enable data alignment.

We pre-registered removal of artefacts >2 seconds based on visual inspection in AcqKnowledge. However, all identified artefacts were <2 seconds and were therefore retained. Data and event markers exported to R studio and downsampled to 500 Hz. SCL values were then z-scored within person and aggregated across the threatening and non-threatening blocks of the virtual world (i.e., separately for the first threatening block, first non-threatening block, second threatening block and second non-threatening block).

Heart rate in the virtual world was recorded using a wireless BIOPAC BioNomadix amplifier (BN-RSPEC), using the same three-lead ECG set-up as the resting heart rate recording. The transmitter belt was attached around participants' chest.

Artefacts >2 seconds were removed from the ECG data based on visual inspection in Acqknowledge (7 artefacts across 5 participants). R-peaks were automatically detected in Acqknowledge and visually checked, as described for the HRV analysis. A list of R-peaks was then exported into R studio and instantaneous heart rate was calculated using the R package *RHRV* (2). Heart rate data were z-scored within person and aggregated across the threatening and non-threatening blocks of the virtual world.

**Methodological Details S3.** Sleep monitoring.

Polysomnography was recorded using an Embla N7000 system (Embla Systems, Broomfield, CO, USA). The scalp was prepared with NuPrep exfoliating agent before gold-plated electrodes were attached at eight standard locations according to the international 10-20 system (3): F3, F4, C3, C4, P3, P4, O1, and O2; each referenced to the contralateral mastoid. Left and right electrooculogram, left, right, and upper electromyogram, and a ground electrode (forehead) were also attached. All electrodes were verified to have a connection impedance of $< 5$ kΩ. All signals were digitally sampled at 200 Hz.

Using RemLogic 3.4, polysomnography data were partitioned into 30 s epochs and scored as wakefulness, non-rapid eye movement sleep N1, N2 or N3 (slow-wave sleep), or rapid eye movement sleep, based on standardised criteria (4). Epochs scored as N2 and N3 were exported to MATLAB 2019a using the FieldTrip toolbox (v10/04/18) for spectral analysis. Noisy channels (identified during sleep scoring) were removed (7 channels across 6 participants) and other artefacts were removed using Fieldtrip's data browser. Artefact-free N2 and N3 epochs across frontal (F3 and F4), central (C3 and C4), and parietal (P3 and P4) channels were applied to a Fast Fourier Transformation with a 10.24 second Hanning window and 50% overlap. Spectral power in the delta band (0.8–4.6 Hz) was divided by absolute power across all frequency bands to produce a normalised index of SWA (5).

**Note S1.** Changes in alertness levels after experimental manipulation.

***Stanford Sleepiness Scale***

To examine whether sleep deprivation reduced alertness levels relative to restful sleep, scores on the Stanford Sleepiness Scale (SSS; 6) were applied to two-way mixed ANOVA with the between-subjects factor *Group* (Sleep Rested or Sleep Deprived) and the within-subjects factor *Session* (Evening or Morning). Whereas SSS scores decreased from evening to morning in the sleep rested group (t = 2.58, $p$ = .016, d = 0.51), they significantly increased in sleep-deprived individuals (t = 7.61, $p$ <.001, d = 1.88; F(1,52) = 59.38, $p$ < .001, $\eta\rho^2$ = 0.53).

We also found significant main effects for both *Group* (F(1,52) = 27.11, $p$ <.001, $\eta\rho^2$ = 0.34) and *Session* (F(1,52) = 22.49, $p$ <.001, $\eta\rho^2$ = 0.30). Sleep-deprived participants reported significantly higher sleepiness in the morning session than sleep-rested participants (t = 8.16, $p$ < .001, d = 2.22). There was no significant between-group difference in the evening session (t = 0.38, $p$ = .706, d = 0.10).

***Psychomotor vigilance task***

To further assess whether sleep deprivation reduced alertness levels compared to a night of sleep, we examined changes in reaction time on a psychomotor vigilance task (7), with mean reaction time scores applied to a two-way mixed ANOVA with the between-subjects factor *Group* (Sleep Rested or Sleep Deprived) and the within-subjects factor *Session* (Evening or Morning). Whereas reaction time did not significantly differ from evening to morning in the sleep-rested participants (t = 1.01, $p$ = .322, d = 0.11), reaction times significantly increased in the sleep-deprived group (t = 4.44, $p$ = <.001, d = 0.98; F(1,52) = 11.94, $p$ = .001, $\eta\rho^2$ = 0.19).

We also found a significant main effect for *Session* (F(1,52) = 19.36, $p$ <.001, $\eta\rho^2$ = 0.27). Sleep-deprived participants showed markedly slower reaction time in the morning session compared to sleep-rested participants (t = 2.58, $p$ = .013, d = 0.70). There was no corresponding between-group difference in the evening session (t = 0.14, $p$ = .887, d = 0.04). There was no main effect for *Condition* (F(1,52) = 2.29, $p$ =.136, $\eta\rho^2$ = 0.04).

**Note S2.** Sleep loss and emotional adaptation to ambiguous threat.

### *Heart rate*

Heart rate was significantly elevated during the second threat exposure (F(1, 52) = 4.83, $p$ = .032, $\eta_p^2$ = 0.09). No significant differences were observed between sleep-deprived and sleep-rested participants (*Group*Block*: F(1,52) = 0.31, $p$ = .580, $\eta_p^2$ < 0.01; *Group:* F(1,52) = 0.01, $p$ = .920, $\eta_p^2$ < 0.01).

### *Self-reported arousal*

Self-reported arousal ratings were significantly elevated during the first threat exposure (F(1,52) = 8.07, $p$ = .006, $\eta_p^2$ = 0.13). No significant differences were observed between sleep-deprived and sleep-rested participants (*Group*Block*: F(1,52) = 0.08, $p$ = .777, $\eta_p^2$ < 0.01; *Group:* F(1,52) = 0.96, $p$ = .331, $\eta_p^2$ = 0.02).

**Note S3.** Sleep loss and resolution of emotional reactivity following ambiguous threat.

### *Heart rate*

Heart rate decreased significantly during the first transition into a non-threatening block ($F(1,52) = 4.25$, $p = .044$, $\eta_p^2 = 0.08$). No significant differences were observed between sleep-deprived and sleep-rested participants (*Group*Block*: $F(1,52) = 0.26$, $p = .610$, $\eta_p^2 = 0.01$; *Group*: $F(1,52) = 0.18$, $p = .677$, $\eta_p^2 < 0.01$).

### *Self-reported arousal*

Changes in self-reported arousal ratings did not significantly differ between the first and second transition into non-threatening blocks ($F(1,52) = 1.03$, $p = .316$, $\eta_p^2 = 0.02$). No significant differences were observed between sleep-deprived and sleep-rested participants (*Group*Block*: $F(1,52) = 0.00$, $p = .950$, $\eta_p^2 < 0.01$; *Group*: $F(1,52) = 0.57$, $p = .453$, $\eta_p^2 = 0.01$).

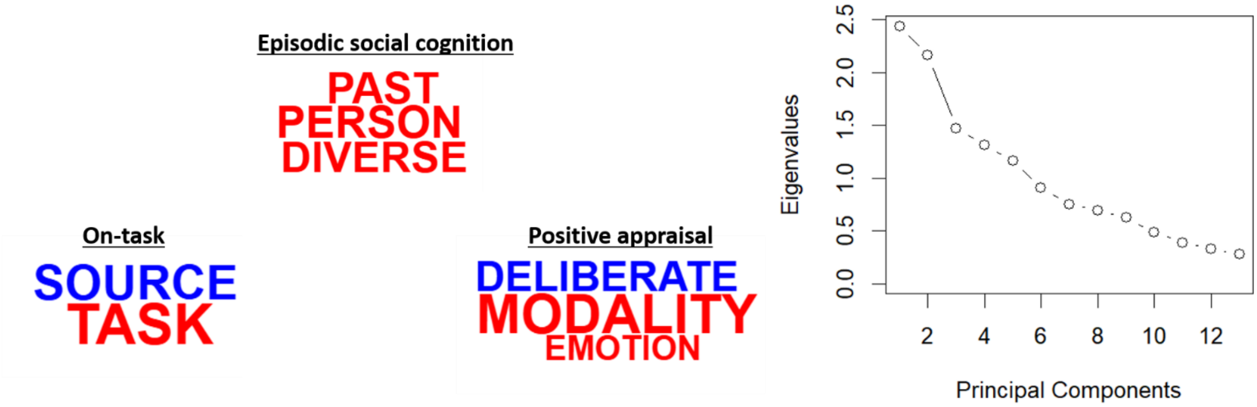


**Figure S1.** Principal component analysis (PCA). Left: three additional components were identified. Component loadings are presented as a word cloud, where the word size reflects loading magnitude and colour indicates direction of the relationship (red: positive, blue: negative). No significant effects emerged when the component loadings were applied to the *Group***Session* ANOVA reported in the main text.
